# The E3 ubiquitin ligase Itch regulates death receptor and cholesterol trafficking to affect TRAIL-mediated apoptosis

**DOI:** 10.1101/2023.05.10.540247

**Authors:** James Holloway, Richard C. Turkington, Daniel B. Longley, Emma Evergren

## Abstract

The activation of apoptosis signalling by TRAIL (TNF-related apoptosis-inducing ligand) through receptor binding is a fundamental mechanism of cell death induction and is often perturbed in cancer cells to enhance their cell survival and treatment resistance. Ubiquitination plays an important role in the regulation of TRAIL-mediated apoptosis, and here we investigate the role of the E3 ubiquitin ligase Itch in TRAIL-mediated apoptosis in oesophageal cancer cells. Knockdown of Itch expression resulted in resistance to TRAIL-induced apoptosis, caspase-8 activation, Bid cleavage and also promoted cisplatin resistance. Whilst the assembly of the death-inducing signalling complex (DISC) at the plasma membrane was not perturbed relative to the control, the TRAIL-R2 receptor was mis-localised in the Itch-knockdown cells. Further, we observed significant mitochondrial widening with an increased cholesterol content. An inhibitor of cholesterol trafficking, U18666A, was able to replicate some of the effects of Itch knockdown, including protection from TRAIL-induced apoptosis, reduced caspase-8 activation, Bid cleavage and Cisplatin resistance. This study highlights the importance of Itch in regulating the crosstalk between mitochondrial cholesterol and TRAIL-induced apoptosis.

## INTRODUCTION

Itch (AIP4) is a member of the of E3 ubiquitin ligase family that mediates lysine-29, -48 and – 63-linked ubiquitin conjugation^1^. It is associated with pro-proliferative activities and endocytic trafficking^2^. Itch is composed of an N-terminal lipid binding C2 domain, a proline rich region, WW motifs and in the C terminus a HECT (homologous to the E6-AP C-terminus) domain. The two WW motifs mediate an intramolecular interaction that keeps Itch in an inactive conformation. A number of substrates have been identified for Itch; including FLIP, Numb, SREBP2, chemokine receptor CXCR4 and Bid^3^. Regulation of the anti-apoptotic protein FLIP is important during cell death, and it has been shown that ubiquitination by Itch promotes its degradation and reduces its protein levels^4, 5^. Itch also interacts with and ubiquitinates a number of endocytic proteins, including endophilin, intersectin-1, amphiphysin, Snx9, Pacsin and Grb2, which results in regulation of trafficking of cell surface receptors^6^. Itch is highly expressed in the gastrointestinal tract, brain and skeletal muscle, and it is overexpressed in multiple types of cancers ^7^.

The regulation of death receptor signalling by ubiquitination has recently emerged as an important mechanism of resistance to cell death in cancer cells ^8, 9^. Core components of the TRAIL signalling pathway have been demonstrated to be regulated by ubiquitination, namely TRAIL-R1/2, FLIP and caspase-8. To evade cell death cancer cells frequently use the ubiquitin-proteasomal system for degradation of pro-apoptotic proteins^10, 11^. TRAIL-mediated apoptosis is unique because it selectively induces apoptosis in cancer cells compared to untransformed cells^12–14^. Binding of the ligand TRAIL (TNF-related apoptosis-inducing ligand) to death receptors (TRAIL-R1, TRAIL-R2) on the cell surface results in recruitment of FADD and activation of procaspase-8 or -10 and the subsequent cleavage of Bid, which results in its translocation to mitochondria and initiation of BAX/BAK pore formation. Bid is a key mediator of both intrinsic and extrinsic apoptosis and has been proposed to define whether TRAIL-mediated apoptosis is mitochondria-dependent or -independent ^15, 16^. A regulator of caspase-8 activity, and Bid cleavage, is the FLICE-like inhibitory protein (FLIP); FLIP is a procaspase-8 homolog that is capable of competing with caspase-8 for binding to FADD and is itself negatively regulated by ubiquitination^3, 5^.

We hypothesized that Itch may play a role in the regulation of TRAIL-induced apoptosis by altering the abundance and/or subcellular distribution of FLIP or TRAIL receptors. In the TCGA pan-cancer atlas the three cancer types with the highest gain of copy number for ITCH were rectum adenocarcinoma (87%, READ), colon adenocarcinoma (69%, COAD) and oesophageal carcinoma (57%, ESCA) making these interesting disease settings in which to study Itch. Consequently, we compared the expression of ITCH in cell lines from these cancers and focussed our work on the cell line with the highest expression of ITCH, OE33, an oesophageal cell line ^17^. In this study, we examined whether knockdown of Itch affects the endocytosis of TRAIL-R2/R1, FLIP expression and TRAIL-mediated signalling in the oesophageal cell line OE33 that displays a high expression of Itch. We demonstrate that TRAIL-mediated cell death and trafficking of the TRAIL-R2 receptor are regulated by the E3 ubiquitin ligase Itch through cholesterol trafficking.

## Materials & Methods

### Cell Lines

OE33, LIM1215 and the KM12 cell lines were maintained in RPMI media (Gibco, UK). COLO320, HCT116 and HT29 cell lines were maintained in Macoy’s5A media (Gibco, UK). Cell culture media was supplemented with 10% foetal bovine serum (Gibco, UK), 5% Penicillin/Streptomycin (Gibco, UK) and 2mM L-glutamine (Gibco, UK). All cells were maintained at 37°C in a 5% CO_2_ humidified atmosphere and were regularly screened for the presence of mycoplasma using the MycoAlert Mycoplasma Detection Kit (Lonza, Switzerland).

### Reagents and Antibodies

The following commercial antibodies were used: TRAIL-R2/DR5 (rabbit, Cat# 3696S, Cell Signalling Technology); BID (rabbit; Cat# 2002S, Cell Signalling Technology); PARP (rabbit, Cat# 9542S, Cell Signalling Technology); Caspase 3 (rabbit, Cat# 9662S, Cell Signalling Technology); FLIP NF6 (mouse, Cat# AG-20B-0056-C100, Adipogen); Itch (mouse, Cat# 611198, BD Biosciences); FADD (mouse, Cat# 556402, BD Pharmigen); Caspase 8 (mouse, Cat# ALX-804-242-C100, Enzo); TRAIL-R1/DR4 (rabbit, Cat# AB16955, Calbiochem); SREBP2 (rabbit, Cat# ab30682, Abcam); beta-Actin (mouse, Cat# A5316, Sigma) and GAPDH (mouse, Cat# sc-47724, Santa Cruz). The following PE-conjugated antibodies were used in cell surface expression experiments: isotype control (Cat# 12-4714-73, eBioscience); DR5 (Cat# 12-9908-42, eBioscience) and DR4 (Cat# 12-6644-42, eBioscience). AMG655 (Conatumumab) was sourced from Amgen Inc (Thousand Oaks, CA, USA); this antibody was coupled to Dynabeads using the Dynabead® Coupling Kit (Life Technologies, Paisley, UK) for use in TRAIL-R2 DISC IP assay. The secondary antibodies used for immunocytochemistry were Alexa Fluor488- and 568-conjugated goat anti-rabbit, anti-mouse and anti-sheep (Thermo Fischer Scientific); for Western blot detection goat anti-rabbit and anti-mouse IgG-HRP conjugates (Cell Signaling Technologies). The following reagents were used: Filipin III (SAE0087; Sigma-Aldrich); DAPI (#62248; Thermo Fischer Scientific); U18666A (#662015; Calbiochem). Human isoleucine-zipper TRAIL (izTRAIL) was expressed and purified as described and stored at -80°C in aliquots^18^.

### Knockdown Cell Line Generation

To generate *ITCH* knockdown cell lines lentivirus expressing short hairpin RNA (shRNA) from the pLKO1.puro plasmid were generated with the targeting sequence of the *ITCH* gene AACACCTCGAGACAACCTC (shEE162) and the shRNA control CAACAAGATGAAGAGCACCAA (Sigma MISSION Target shRNA). Selection was carried out using 1µg/ml of puromycin for 48 hours. The efficiency of shRNA knockdown was assessed by Western blot.

### siRNA Transfections

All siRNAs were obtained from Dharmacon (Chicago, IL, USA), and transfections were carried out using Lipofectamine RNA iMAX (Life Technologies, Paisley, UK) as previously described. The smartpool sequences for ITCH were; GUUGGGAACUGCUGCAUUA, CAACAUGGGACGUAUUUAU, GAAAUUAAGAGUCAUGAUC and CGAAGACGUUUGUGGGUGA.

### Immunoblotting

Whole cell lysate was prepared in RIPA buffer and Western blot analysis was carried out as described previously^19^. Protein expression was detected using the Western Lighting Plus-ECL substrate from PerkinElmer (Waltham, MA) on a G:BOX Chemi XX6 gel doc system (Syngene). Densitometry was carried out using ImageJ software.

### AnnexinV/PI Staining

Samples were analysed on a BD Accuri C6 Plus flow cytometer (BD Biosciences). Initially, 2x10^5^ cells were seeded into wells of a 6-well plate in duplicate, left 24 hours before addition of drug and cell death was assessed following live cell staining with FITC-tagged AnnexinV (BD Biosciences) and addition of PI (Sigma) directly before analysis.

### Cell surface flow cytometry

Cell surface TRAIL-R1 and TRAIL-R2 expression was assessed following live cell staining with Phycoerythrin-conjugated antibodies on a BD Accuri C6 Plus flow cytometer. All experiments were gated using an isotype control antibody. 10,000 cells were analysed per sample in duplicate. Experiments were repeated three times.

### Cell viability assay

Cell viability was assessed using CellTiterGlo Luminescent Assay (Promega, Madison, WI) according to manufacturer’s instructions. Briefly, 4x10^3^ cells per well were seeded into a white opaque 96-well plate. Each sample was plated in triplicate. Drugs were added in a total volume of 200µl. Once experimental procedures were met, 150µl of media was removed and 50µl of Glo reagent was added to the wells. The plate was left to agitate gently at room temperature for 10 minutes before analysis on a Biotek plate reader was carried out using a 1 second luminescence read.

### Caspase activity assay

Caspase activity was assessed using Caspase-3/-7 Glo Luminescent Assay (Promega, Madison, WI) according to manufacturer’s instructions. Briefly, 4x10^3^ cells per well were seeded into a white opaque 96-well plate. Each sample was plated in triplicate. Drugs were added in a total volume of 50µl per well. Once experimental procedures were met, 50µl of appropriate Caspase Glo® Luminescent Assay was added to each well. The plate was left to agitate gently at room temperature for 45 minutes before analysis on a Biotek plate reader was carried out using a 1 second luminescence read.

### DISC-IP

The TRAIL-R2 DISC-IP was carried out as previously described^20^. Briefly, 30μl of Dynabeads (Thermo Fisher, UK) coated with 2.5μg of AMG655 was incubated with the cells for 30 minutes. Cells were lysed with DISC IP buffer (0.2% NP-40, 20 mM Tris–HCL (pH 7.4), 150 mM NaCl and 10% glycerol) for 60 min on an orbital shaker at 4°C. The beads were subsequently washed five times in DISC IP buffer before recruitment of DISC proteins was assessed by Western blotting.

### Immunofluorescence

Immunocytochemistry was performed as described previously^19^. Briefly, cells were plated onto 13mm glass coverslips, fixed at room temperature in 4% paraformaldehyde in phosphate buffered saline (PBS). To visualise the cell surface with Alexa488-tagged wheat germ agglutin (Thermo Fisher Scientific, UK, 1:2000 dilution) an incubation was done prior to permeabilization. Cells were stained with primary and secondary antibodies and mounted on glass slides. Confocal images were acquired of the cells at room temperature using a Leica SP8 confocal microscope equipped with a 63x objective (1.4 NA HCX PL APO lens) controlled by the Leica Application Suite-X software. For each experiment 20 cells were analysed.

### Image analysis

Confocal image analysis was carried out using the ImageJ® software (Fiji). Receptor expression was expressed as the Integrated Density of the stain at the plasma membrane divided by the area. The plasma membrane was identified using wheat germ agglutin stain and the integrated density was measured by highlighting the intensity of the receptor staining at the plasma membrane. Colocalization analysis was performed using the Pearson colocalization coefficient from 14 cells per group using the JaCOP plugin in ImageJ.

### Transmission electron microscopy

Embedding and processing of samples was performed as described^19^. Images were acquired on a Jeol JEM1400plus microscope at 80kV equipped with a JEOL Ruby (8 MPixel) bottom-mounted CCD camera.

### Statistical analysis

Statistical significance was calculated from distinct technical replicates (nL≥L3; unless otherwise stated), either by Student’s T test (two-tailed) or two-way ANOVA in GraphPad Prism 9. Two-way ANOVA tests were performed to compare two factors (for e.g. shRNA and drug concentration). Multiple comparison analyses were performed using GraphPad Prism 9. Graphs were plotted as means with error bars represented as SEM; statistical significance was denoted as follows: ***pL<L0.001, **pL<L0.01, *pL<L0.05, nsL=LpL>L0.05. Experimental phenotypes were confirmed in at least three independent experiments.

## RESULTS

### Itch regulates TRAIL-mediated apoptosis independently of FLIP

To investigate the role of Itch in TRAIL-mediated apoptosis, we generated a stable knockdown in the oesophageal cancer cell line OE33 and subjected it to TRAIL treatment followed by quantification of cell viability and apoptosis. Western blot analysis demonstrated a reduction in protein expression of 80% in the OE33 Itch knockdown (KD) cell line with no change in the expression of pro-caspase-8 or the two FLIP splice forms (**Figure 1A-C**). A similar lack of impact on FLIP expression was observed in a panel of oesophageal and colorectal cancer cell lines (**Supplementary Figure 1**). FLIP(L) has been shown to be ubiquitinated by Itch in a few cancer cell lines and fibroblasts, while in macrophages its stability is not regulated directly by FLIP(L)^5, 21–24^.

**Figure 1:**
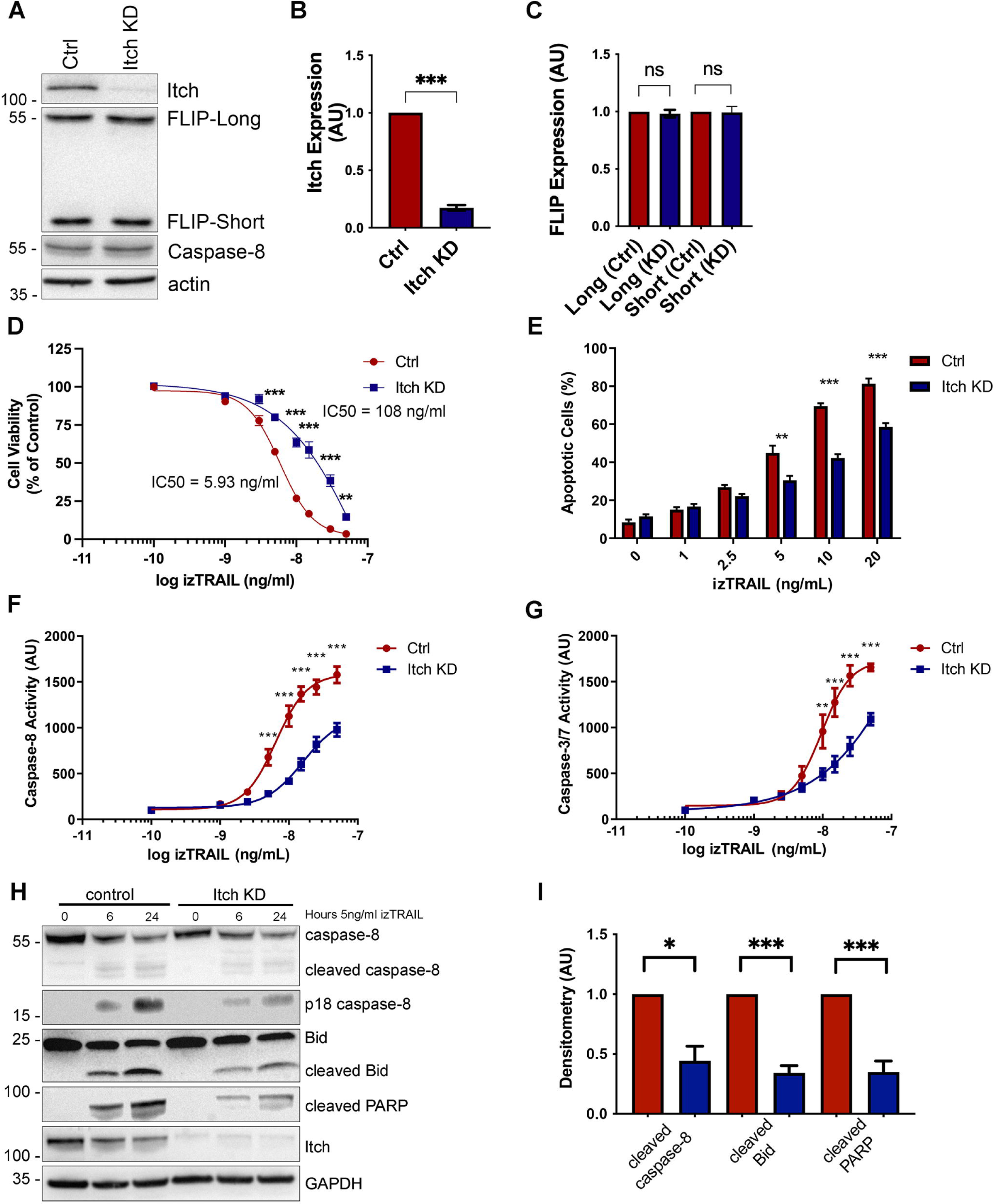
Itch knockdown increases resistance to TRAIL mediated apoptosis in OE33 cells and impairs caspase-8 cleavage and activity. **(A)** Western blot analysis of basal expression of FLIP(L), FLIP(S), FADD, Procaspase-8, TRAIL-R1, TRAIL-R2 in OE33 Ctrl (shCTRL) and Itch knockdown (KD) cells stably expressing shRNA targeting ITCH. **(B)** Bar graph showing the densitometry of Western blot analysis of Itch expression normalised to the actin loading control in five independent experiments. **(C)** Bar graph showing the quantification fo FLIP-Long and -Short expression in control shRNA and Itch knockdown (KD) cell lines. N=5 independent experiments. **(D)** OE33 Ctrl and Itch KD cell lines were subjected to treatment with increasing doses of izTRAIL for 24h (0-50ng/mL). Cell viability was measured using the CellTiterGlo® Assay. Data was normalised to an untreated control. **(E)** OE33 Ctrl and Itch KD cells were treated with 0-20ng/mL TRAIL for 24 hours and then stained with FITC-conjugated Annexin V and propidium iodide to assess the number of apoptotic cells under each treatment condition. Bar graph showing the percentage of apoptotic cells (Annexin V-positive cells) of the population. **(F)** Caspase -8 and **(G)** Caspase -3/-7 activity in OE33 Ctrl and Itch KD cell lines was measured following treatment with increasing concentrations of izTRAIL (0-50ng/mL). Activity was measured 6 hours post treatment using the Caspase -3/-7 or the Caspase -8 Glo® assays. **(H)** Western blot analysis of cleaved Caspase-8, Bid and PARP in OE33 Ctrl and Itch KD cell lines following treatment for 6 or 24 hours with 5ng/ml izTRAIL. **(I)** Densitometry of the experiment in C in four independent repeats. Error bars represent the standard error of the mean. Statistical significance was measured by Student’s t-test. *, p <0.05, ** p< 0.01, ***, p<0.001.

To assess the functional effects of Itch knockdown on TRAIL-induced apoptosis, we used recombinant isoleucine-zipper (iz) TRAIL, which maintains the trimeric nature of the cell surface TRAIL ligand and enhances its activity ^25^. The Itch KD cell line was significantly more resistant to the izTRAIL-induced apoptosis as assessed by cell viability (p<0.001; **Figure 1D**) and apoptosis assays (p<0.001; **Figure 1E**). The Itch KD cell line also displayed significantly reduced activity of caspase-8 (p<0.001; **Figure 1F**) and caspase-3/7 (p<0.001; **Figure 1G**) compared to the control cell line following treatment with izTRAIL. These results were corroborated by Western blot analysis that showed a reduction of cleaved caspase-8, PARP and Bid in the Itch KD cell line (**Figure 1H,GI;** p<0.001). In conclusion, our findings demonstrate that Itch regulates TRAIL-mediated apoptosis through a mechanism that is not dependent on FLIP.

### Itch knockdown causes a mis-localization of TRAIL-R2 in OE33 cells

Next, we sought to investigate whether loss of Itch mediates TRAIL resistance through downregulation of the cell surface expression of death receptors. A major limiting factor of apoptotic TRAIL signalling is the availability of the receptor at the cell surface^26, 27^. Itch targets components of the endocytic machinery and has been shown to regulate signalling and stability of cell surface receptors^28^. A significant reduction in cell surface TRAIL-R2 expression was observed in the Itch knockdown compared to the control (**Figure 2A**; p<0.001). In comparison, the cell surface expression of TRAIL-R1 (DR4) was found to be unaltered between the two cell lines (**Figure 2B**), while Epidermal Growth Factor receptor (EGFR) cell surface staining was increased (p<0.05; **Figure 2C**) and transferrin internalisation was unchanged (**Figure 2D**). TRAIL-R1, -R2 and EGFR are internalised after ligand binding by clathrin-dependent and -independent endocytosis while the transferrin receptor is dependent on clathrin-mediated endocytosis ^29^. Therefore, these data suggested that clathrin-mediated receptor endocytosis was not enhanced by Itch depletion and altered receptor trafficking could not explain the reduction of TRAIL-R2 on the cell surface. Immunocytochemistry supported the finding that TRAIL-R2 cell surface expression was reduced in the Itch KD OE33 cell line (**Figure 2E**). Importantly, a loss of Itch expression did not have an impact on total TRAIL-R1/R2 receptor (DR4/5) protein expression in cell lysates (**Figure 2F**). To validate the findings further, a panel of oesophageal and colorectal cancer cell lines were used which also displayed a reduced TRAIL-R2 cell surface expression upon Itch knockdown (**Supplementary Figure 2**). In summary, knockdown of Itch resulted in a resistance to TRAIL-mediated apoptosis and correlated with a loss of TRAIL-R2 at the plasma membrane but not TRAIL-R1. This indicated that Itch either regulates anterograde membrane trafficking of TRAIL-R2 (but not TRAIL-R1), enhances the assembly of the DISC complex, or regulates caspase-8 and/or Bid association with mitochondria. The first option seemed unlikely as there are no data suggesting that the two TRAIL receptors traffic to the plasma membrane via independent mechanisms and we therefore focussed on exploring the other two options.

**Figure 2:**
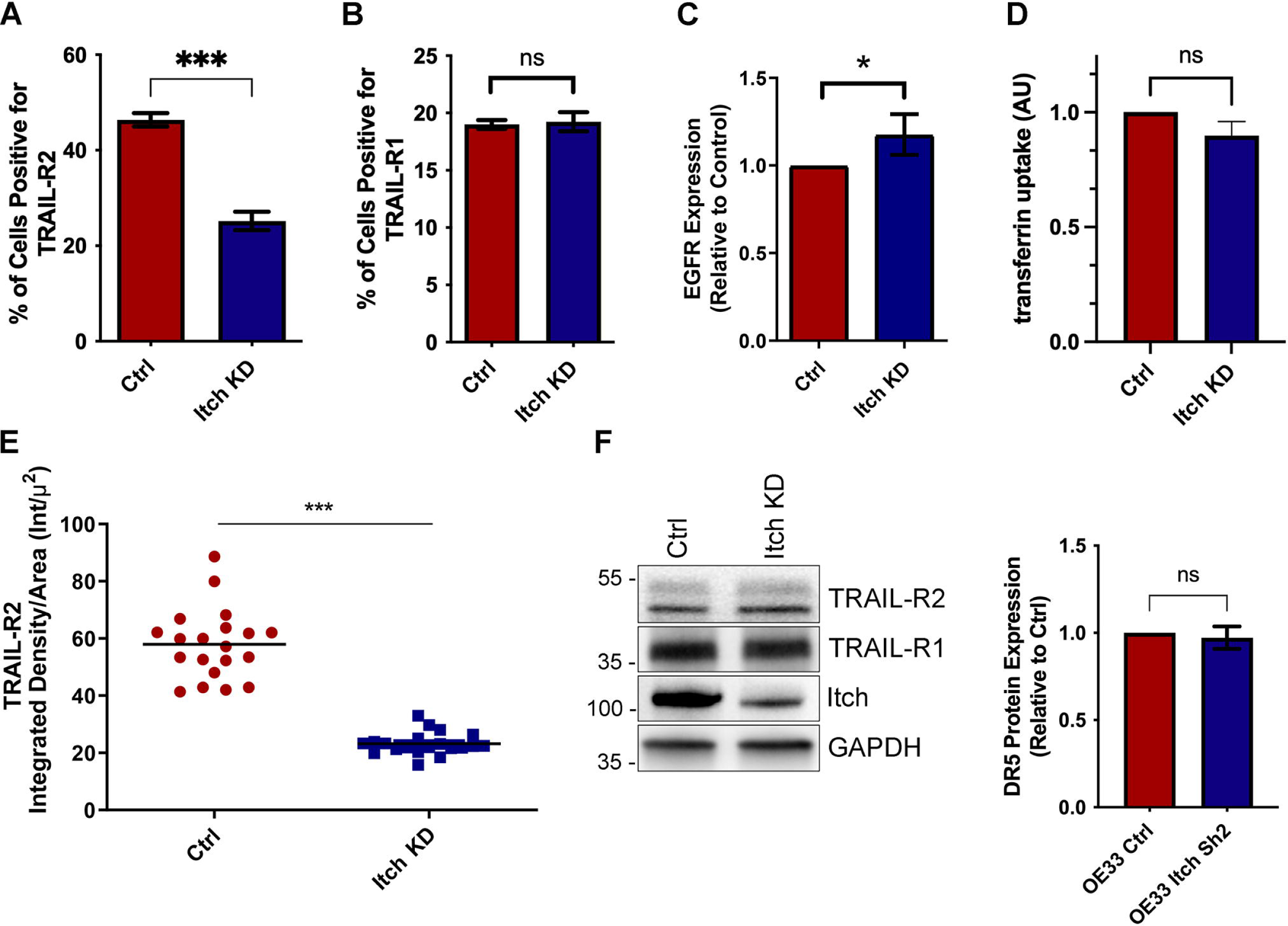
Itch knockdown cells show reduced surface expression of TRAIL-R2 in OE33 cells. FACS analysis of cell surface expression of **(A)** TRAIL-R2 and **(B)** TRAIL-R1 cell surface expression in OE33 Ctrl and Itch KD cell lines. The fluorescence intensity of cell surface staining is represented as percentage of receptor positive cells compared to an IgG isotype control. **(C)** FACS analysis of cell surface expression of EGFR in serum starved cells normalised to the control cell line in three independent experiments. **(D)** FACS analysis of fluorescent transferrin uptake in serum starved cells. Three independent experiments were performed where 10,000 cells were analysed per experiment. **(E)** TRAIL-R2 cell surface expression was measured using confocal microscopy. Alexa488-tagged wheat germ agglutinin stain was used to identify the plasma membrane and Anti-DR5 stain was used to determine the intensity of TRAIL-R2 at the plasma membrane. Receptor expression is expressed as the Integrated Density of the stain at the plasma membrane divided by the area. **(F)** Western blot showing the total expression levels of TRAIL-R2 and -R1 in control and Itch KD OE33 cells. Error bars represent the standard error of the mean. Statistical significance was measured by Student’s t-test. *, p <0.05, ***, p<0.001.

### TRAIL-activated DISC assembly is independent of Itch

We hypothesised that Itch may be incorporated into the death receptor signalling complex (DISC) where it could regulate caspase-8 recruitment or activation, and therefore next investigated whether Itch regulates the assembly of the DISC, which is induced by TRAIL. A reduction in TRAIL-R2 recruitment into the DISC in the Itch KD cells was observed (**Figure 3A**), consistent with reduced cell surface levels of TRAIL-R2 (**Figure 2A**). However, this did not correlate with significantly reduced recruitment of caspase-8, FADD or FLIP(L). Furthermore, caspase-8 dependent cleavage of FLIP-L at the DISC was maintained in the Itch KD cell line, and all other components of the complex are there in equal amounts to the control suggesting that Itch does not regulate the DISC assembly or activation of caspase-8 at the DISC. Taken together, our findings indicate that Itch regulation of caspase-8 activity and apoptosis is not directly due to alterations in the death receptor complex at the plasma membrane.

**Figure 3:**
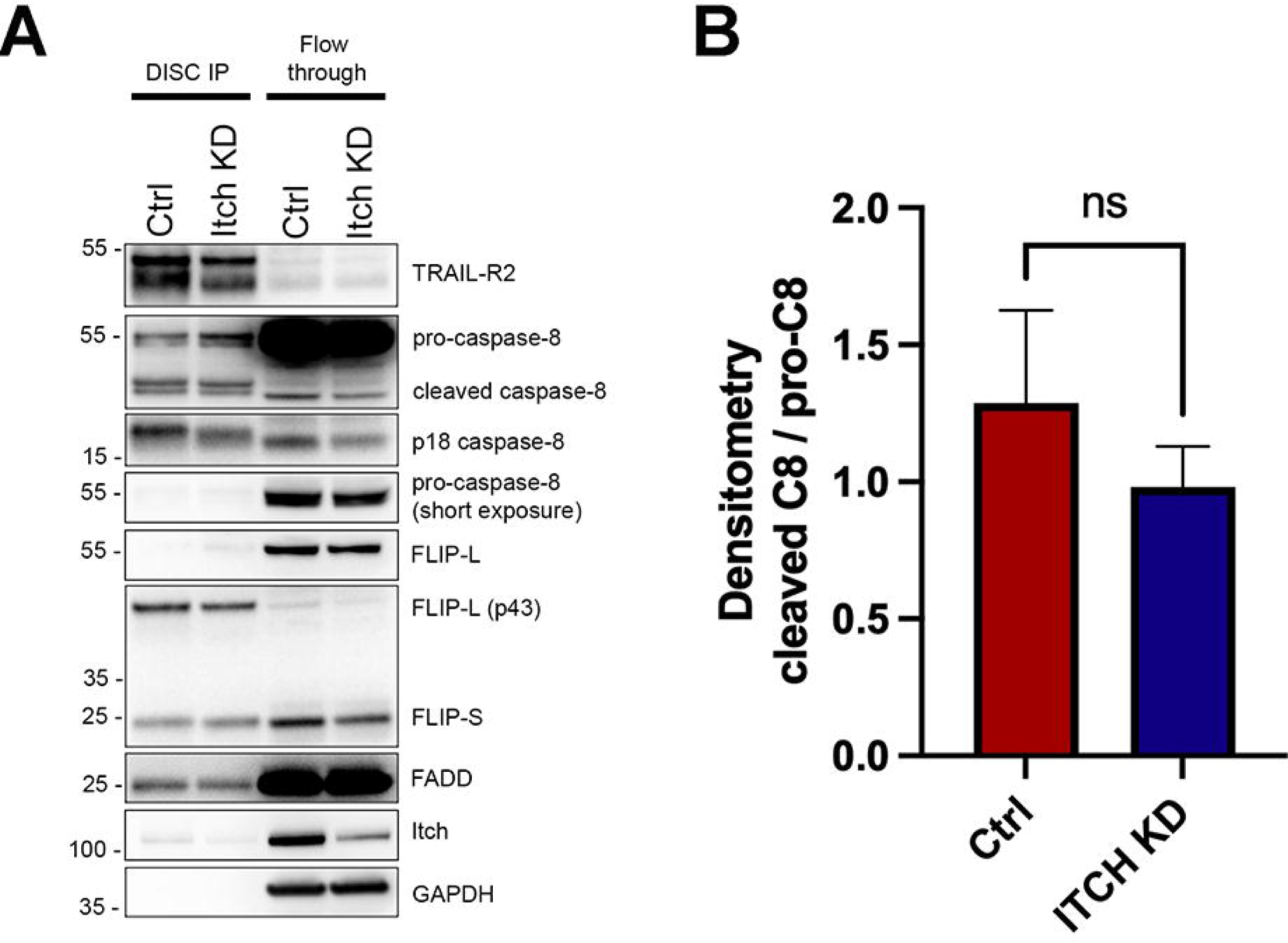
Itch is found at the death-inducing signalling complex (DISC) but does not regulate its composition or cleavage of caspase-8. **(A)** Immunoprecipitation of the DISC using an activating TRAIL-R2 antibody. Cells were treated for 1 hour to induce formation of the DISC. Samples from the control and Itch KD cell lines were loaded in duplicates and proteins were separated by size on a SDS-PAGE gel prior to Western blot analysis.

### Itch knockdown regulates mitochondrial cholesterol content which implicates a novel regulatory role in apoptosis

On the basis of our findings showing that in the Itch KD cell line receptor endocytosis was not significantly altered and that the assembly of the DISC and recruitment of Caspase-8 is not impaired we next investigated the role of mitochondria in apoptosis in these cells. The OE33 cell line is classified as a type II cell that relies on the intrinsic apoptotic pathway. In type II cells, the lipid composition of the mitochondrial outer membrane is an important regulator of caspase-8 recruitment and activity ^30, 31^. Caspase-8 that is localised to mitochondria interacts with and cleaves Bid to induce MOMP resulting in a release of cytochrome c ^32^. Furthermore, mitochondrial cholesterol mitigates against apoptosis by increasing the stiffness of the mitochondrial membrane to prevent perforations and release of cytochrome c^31^. Experiments in the Itch knockout mouse showed that it regulates cholesterol metabolism through ubiquitination of nuclear SREBP2 and SIRT6 ^39^ resulting in increased intracellular cholesterol levels ^33^. We therefore investigated whether a defect in cholesterol metabolism in the Itch KD OE33 cell line could affect mitochondrial cholesterol content, caspase-8 activity and cell viability^33^.

To investigate the morphology of intracellular organelles, OE33 cells were grown in normal media and imaged by transmission electron microscopy. While the morphology of the ER and Golgi appeared normal, the morphology of the mitochondria was significantly altered (**Figure 4A-B**). Control mitochondria had a normal oblong shape, while mitochondria in the knockdown cells were often round and significantly wider (p<0.001; **Figure 4C**). The mitochondria in the knockdown presented with disorganised cristae, a phenotype similar to that observed in cells where cholesterol accumulate in mitochondria ^34^. To investigate whether there is an increase in mitochondrial cholesterol cells were stained with Filipin-III dye (**Figure 4D**). The mitochondrial outer membrane protein TOMM20 was used as a mitochondrial marker. A prominent Filipin-III stain was observed in mitochondria of Itch KD cells. Quantification of the stain showed a significant increase in colocalization of mitochondria with the Filipin-cholesterol stain in the KD compared to control (p<0.0001; **Figure 4E**). The images further illustrate the altered morphology of mitochondria in cells with reduced Itch expression, where the control mitochondria have a classical oblong shape, whereas in the knockdown the mitochondria appear enlarged. Finally, we investigated whether SREBP2 expression was altered in the OE33 knockdown cell line as it is a main regulator of cholesterol metabolism. When cholesterol levels are low SREBP2 is proteolytically activated and translocated to the nucleus to regulate transcription of genes involved in cholesterol synthesis and uptake. A reduction in the mature nuclear form of SREBP2 was observed in the knockdown compared to the control indirectly indicating elevated cholesterol levels (**Figure 4F**). In summary, the data show that Itch regulates cholesterol content in mitochondria and reduces the expression of the processed form of SREBP-2, both of which indicate that loss of Itch promotes cholesterol metabolism and trafficking.

**Figure 4:**
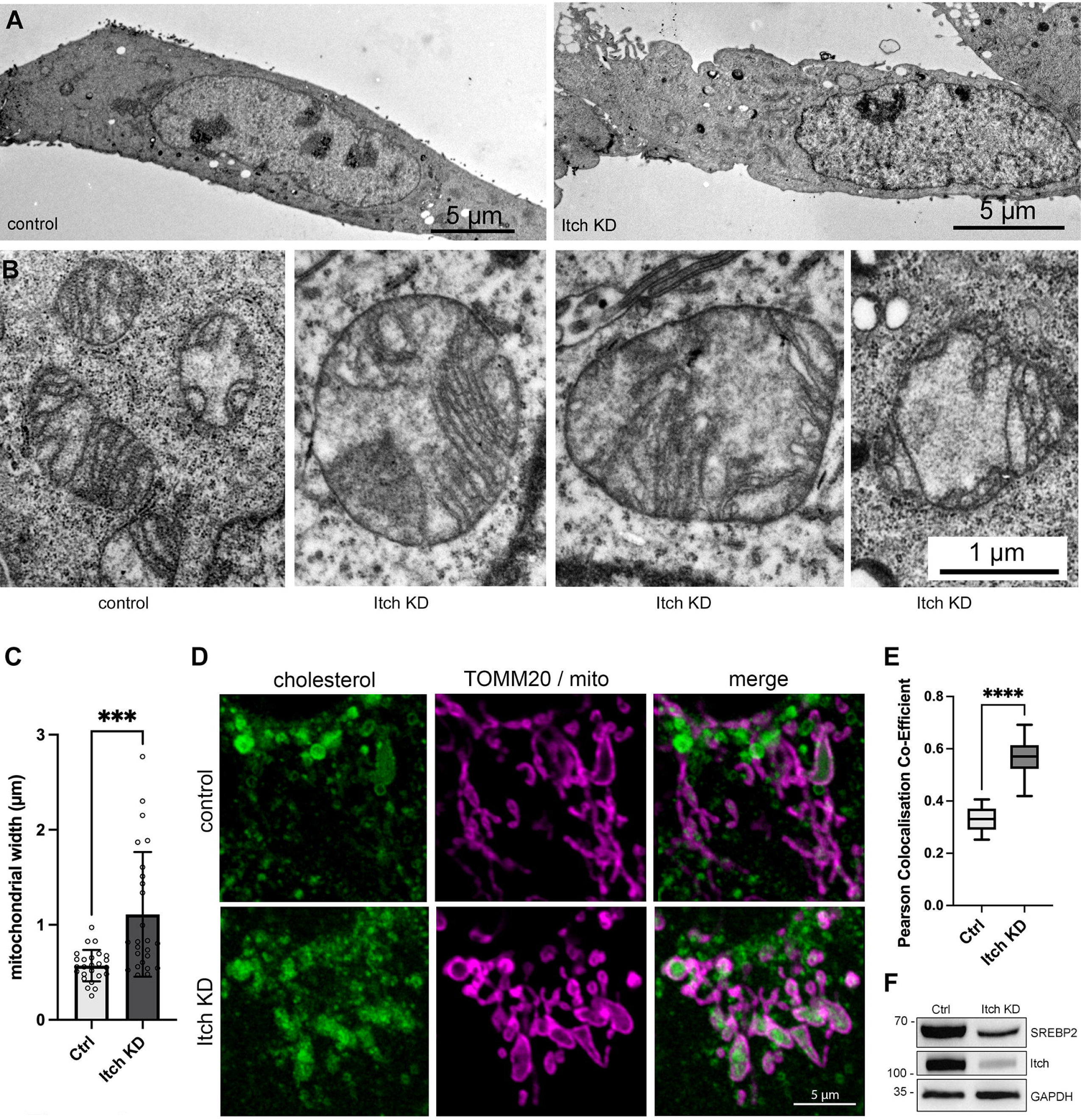
Itch knockdown leads to an accumulation of mitochondrial cholesterol. **(A)** Transmission electron micrographs illustrating the morphology of control and Itch KD cells. Scale bars 5 µm. **(B)** Electron micrographs illustrating the morphology and size of mitochondria in control and Itch KD cells. Scale bar = 1µm for all images. **(C)** Quantification of mitochondrial width from 25 cells. Mean ± SD. **(D)** Confocal images illustrating the accumulation of free cholesterol by Filipin staining together with mitochondrial marker TOMM20. Scale bar, 5µm. **(E)** Tukey box plot displaying the Pearson colocalization co-efficient of TOMM20 and Filipin staining to compare levels of mitochondrial cholesterol in control and Itch KD cell lines. N=14 per group. **(F)** Western blot analysis of cell lysates showing the expression of nuclear SREBP2 (cleaved) protein in the Itch KD compared to control.

### Accumulation of cholesterol in mitochondria cause resistance to TRAIL treatment and inhibits caspase-8 cleavage

Based on our findings that Itch knockdown results in an accumulation of cholesterol in mitochondria we hypothesised that this altered cholesterol trafficking was the underlying cause of the resistance to TRAIL-mediated apoptosis in the Itch knockdown cells. To test this hypothesis, control cells were treated with the drug U18666A to promote trafficking and accumulation of cholesterol in mitochondria^31^. TRAIL-induced activation of Caspase-8 and Caspase-3/7 in control cells treated with U18666A was inhibited to levels similar to that of Itch knockdown cells (**Figure 5A-B**). U18666A used as a single agent did not have any effect on Caspase-8, -3 or PARP cleavage (**Figure 5C**, lanes 1-2), but in combination with TRAIL inhibited activation of Caspase-8, -3 and cleavage of Bid and PARP (**Figure 5C**, lanes 3-4). The expression of TRAIL-R2 or FLIP were not altered by the treatment. Furthermore, treatment with U18666A reduced the cell surface expression of TRAIL-R2 (**Figure 5D**) and increased cholesterol in mitochondria (**Figure 5E**), which aligns with the Itch knockdown data. Together, these experiments provide evidence for a regulatory role for Itch in regulation of caspase-8 activation through mitochondrial cholesterol content and suggest that Itch mediates TRAIL resistance mainly through regulation of cholesterol trafficking. Because cleaved Bid synergises and primes cells for apoptosis upon cisplatin treatment, we hypothesised that OE33 Itch KD cells would be resistant to cisplatin treatment due to the reduced Bid levels^35^. Cell viability assays demonstrated that the Itch KD cell line was significantly more resistant to cisplatin-mediated apoptosis (p<0.001; **Figure 5F**). A three-fold increase in IC_50_ value in the Itch KD cell line for cisplatin (2.75µg/ml) compared to the control cell line (0.91µg/ml) was observed. A significantly reduction of apoptotic cells in the Itch KD was observed at the highest doses of cisplatin (**Figure 5G**). To investigate the impact of mitochondrial cholesterol on cisplatin-induced apoptosis OE33 cells were treated with UA18666A followed by cisplatin (**Figure 5H**), which resulted in cleavage of caspase-8, – 3, Bid and PARP (lanes 1-3). Pre-treatment of cells with U18666A largely abrogated cleavage of apoptotic markers (lanes 4-6). In summary, the data supports a mechanism where Itch and mitochondrial cholesterol trafficking has and important regulatory role in TRAIL- and cisplatin-induced apoptosis.

**Figure 5:**
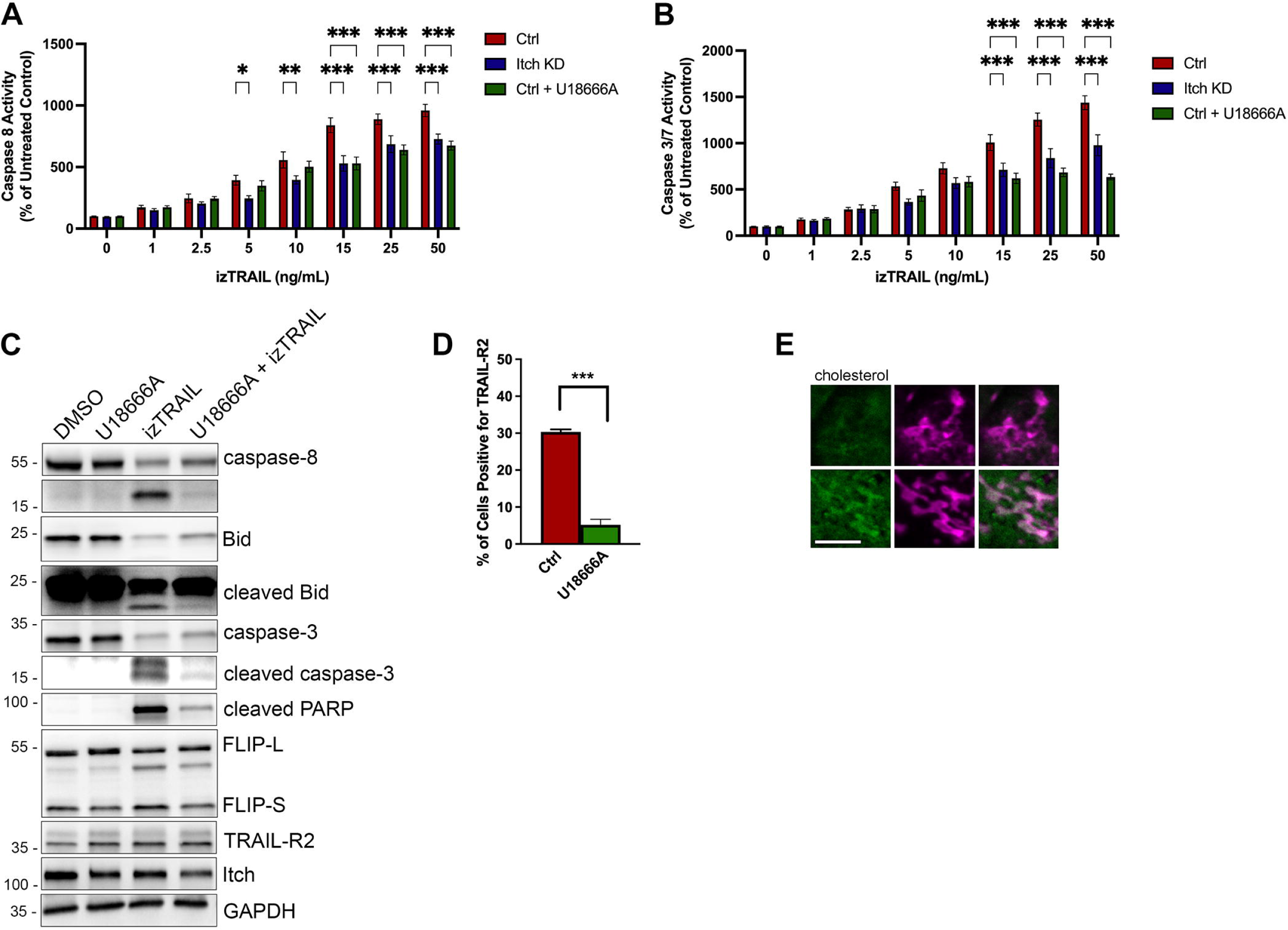
Cholesterol homeostasis is an important resistance mechanism for TRAIL-induced apoptosis. **(A)** Caspase-8 activity and **(B)** Caspase -3/-7 activity was measured following treatment with increasing concentrations of izTRAIL (0-50ng/mL). Activity was measured 6 hours post treatment using the Caspase -3/-7 or the Caspase -8 Glo® assays. Cells were treated with U18666A (1ng/ml) for 24 hours. **(C)** Western blot analysis of cell lysates after treatment with U18666A and izTRAIL for key apoptotic proteins; caspase-8, -3, Bid and PARP. **(D)** Bar graph showing the cell surface expression of TRAIL-R2 in control and cells treated with UA18886 in three independent experiments. **(E)** Fluorescence images showing the accumulation of cholesterol, stained with FilipinIII, in mitochondria after U18666A treatment. Scale bar, 5µm. **(F)** Cell viability assay of the Itch KD cell line in response to Cisplatin treatment for 72 hours at 0-6 µg/ml. **(G)** Apoptosis assay using the Itch KD cell line treated with 0-6 µg/ml Cisplatin for 72 hours. Data was expressed as a percentage of control. (**H**) Analysis of apoptotic markers in whole cell lysates from cells treated with Cisplatin in U18666A. (**I**) A schematic illustrating the proposed role of Itch in regulation of apoptosis.

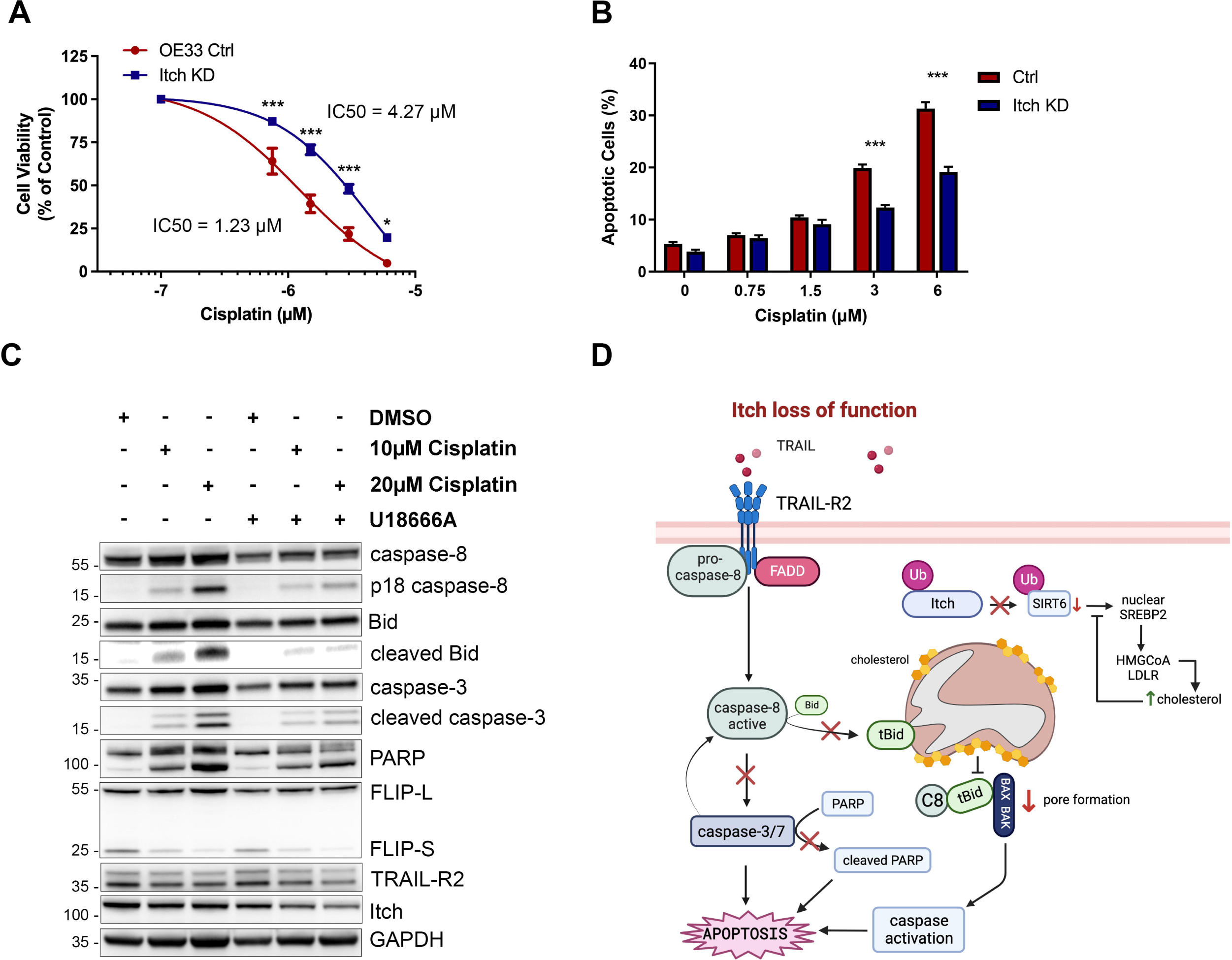

## Discussion

Ubiquitination and deubiquitination have emerged as important regulators of death receptor signalling. Particularly in cancer cells the downregulation of pro-apoptotic proteins by ubiquitination have been shown to mediate resistance to cell death. For example, the ubiquitin ligases Cbl and Cbl-b have been associated with TRAIL-mediated apoptosis resistance mechanisms at the plasma membrane where they regulate receptor clustering in lipid rafts and caspase-8 activation^36, 37^. The fact that these ubiquitin ligases regulate both receptor trafficking to cholesterol-rich lipid domains and downstream TRAIL signalling implies that there may be additional regulatory mechanisms that affect TRAIL sensitivity. Here we have investigated the function of the ubiquitin ligase Itch in this context. Itch is known to regulate receptor trafficking, intracellular cholesterol synthesis, the stability of the anti-apoptotic protein FLIP and the pro-apoptotic protein tBid^3, 28, 33, 38^. Here, we show for the first time that the ubiquitin ligase Itch regulates TRAIL-mediated intrinsic apoptosis through regulation of caspase-8 activation and intracellular cholesterol homeostasis.

In this study, we observed that Itch knockdown OE33 cells are more resistant to TRAIL-mediated apoptosis compared to the control. This correlated with a reduced presence of TRAIL-R2 at the plasma membrane rather than the expected increased expression of the anti-apoptotic protein c-FLIP. The expression levels of TRAIL-R2 protein levels were not impaired and the localisation of TRAIL-R1 or EGFR were not affected. This suggested that it was not due to Itch-regulated receptor endocytosis that was promoting trafficking of the receptors to intracellular stores. Further supporting this notion, clathrin-mediated endocytosis is closely regulated and only two proteins, HIP1 and FCHo2, have been found to increase receptor internalisation ^39, 40^. The intracellular signalling through caspase-8 and caspase-3/7 was found to be reduced in the cell lines with reduced Itch expression, but caspase-8 cleavage at the DISC was not reduced which indicated that the regulation of mitochondrial-dependent apoptosis was contributing to the phenotype. Furthermore, we observed a significant reduction of t-Bid, which after activation by caspase-8 translocates to mitochondria where it associates with Bax and Bak and induces membrane pore formation and subsequent cytochrome c release^41^. The sensitivity to TRAIL signalling is regulated by the targeting of t-Bid by the ubiquitin/proteasome system and the mitochondrial membrane composition^3, 31, 42, 43^. A significant accumulation of cholesterol was observed in mitochondria in the Itch KD, along with reduced Bid and PARP cleavage. This phenotype was reproduced with the drug U18666A that promotes cholesterol trafficking and accumulation of cholesterol in mitochondria. This study provides new insights into the complexity of the intersection between receptor signalling, membrane trafficking and the significance of the ubiquitin E3 ligase Itch in regulating mechanisms of TRAIL-mediated apoptosis.

An important pathway for trafficking of cholesterol from lysosomes to mitochondria occurs through membrane contact sites, and this can be induced by treatment with the drug U18666A^44^. An increase in mitochondrial cholesterol has a strong anti-apoptotic activity due to the inhibition of BAX and BAK, which prevents release of cytochrome c and induction of apoptosis^31^. Thus, an important anti-apoptotic regulator is the lipid composition of not only the plasma membrane but also the mitochondrial outer membrane. Recently it was shown that the lipid composition of the mitochondrial outer membrane has an important role in the recruitment and activation of caspase-8 and subsequent cleavage of Bid^30, 32, 45^. In our experiments we treated cells with U18666A, and observed a loss of cell viability, caspase-8 activity and cleavage of downstream targets similar to that observed in Itch KD cells. Research has shown that inhibition of intracellular cholesterol trafficking by the drug U18666A in the HCT116 cell line leads to a mislocalisation of TRAIL-R2 from the plasma membrane to intracellular organelles, loss of caspase-8 activation, PARP cleavage and 5-FU resistance^46^.

Reprogrammed lipid metabolism is an established hallmark of cancer, and in particular upregulation of the mevalonate pathway. Cholesterol promotes cell proliferation and is regulated by oncogenes (myc, PTEN) and the tumour suppressor p53. Moreover, high mitochondrial cholesterol and its import is reported to be linked specifically with cancer cell resistance to apoptosis and chemotherapy^47–49^. SREBP2 is the transcriptional master regulator of the mevalonate pathway, and it is upregulated in a number of tumour types including oesophageal and colorectal cancer^50, 51^. Accumulation of mitochondrial cholesterol in hepatocellular carcinoma, HeLa and colon cancer cells impairs Bax oligomerisation and mitochondrial-dependent apoptosis^47, 52, 53^. The accumulation of cholesterol has been correlated with a reduction in either a cholesterol exporter or a stabilisation of an importer, resulting in increased mitochondrial cholesterol^47, 54^. The mechanism is due to a stiffening of the membrane due to cholesterol incorporation and an impaired Bax oligomerisation, resulting in apoptosis resistance^31, 47, 53^. Here we propose that Itch-mediated poly-ubiquitination regulates proteins facilitating mitochondrial cholesterol import or export.

Chemotherapy resistance is a common issue in cancer therapy that is associated with cholesterol regulation and in particular cholesterol content in the plasma membrane and mitochondria ^31, 55–59^. Changes in intracellular cholesterol metabolism in cancer cells is a resistance mechanism for chemotherapy and radiation therapy in addition to promoting cell proliferation, invasion and metastasis^60–62^. Our data shows that an increase in mitochondrial cholesterol due to loss of Itch expression or treatment with U18666A correlates with Cisplatin resistance. Our data suggests that it is important to understand both the localisation and role of cholesterol in TRAIL-mediated apoptosis and chemotherapy resistance.

Here we report a novel regulatory mechanism for the ubiquitinating enzyme Itch in TRAIL-mediated apoptosis (**Figure 5I**). One important implication of this work is that the substrates of Itch that are linked to the cholesterol metabolism and trafficking are regulators apoptosis. It will be important to further dissect these mechanisms to map the interrelationship between Itch, TRAIL-R2, cholesterol, and cisplatin resistance. Furthermore, our data suggest that development of biomarkers based on this mechanism may facilitate the selection of patients with TRAIL-responsive tumours.

## Acknowledgements

We thank Prof. Henning Walzcak (UCL Cancer Institute, UK) for generously sharing the bacterial expression construct for isoleucine zipper TRAIL. This work was supported by Northern Ireland Department for the Economy (EE).

## Author contributions

E Evergren, DB Longley and J Holloway conceived and planned the experiments. J Holloway performed the cell culture, flow cytometry sorting and Western blots. J Holloway prepared samples and performed confocal imaging. E Evergren prepared samples and performed transmission electron microscopy. J Holloway performed all other experiments and analysed the data. R Turkington provided expertise and oesophageal cell lines. E Evergren, DB Longley and J Holloway wrote the manuscript.

## Ethics statement

Ethics approval was not required for this work.

## Figure Legends

**Supplementary Figure 1: Investigation of FLIP expression in a panel of cell lines with Itch knockdown.**

Western blot analysis of basal expression of Itch, FLIP(L) and FLIP(S) in cell lines with a stable shRNA-mediated Itch knockdown (**A**) Lim-1215, (**B**) Colo 320, (**C**) KM12, or a transient siRNA-mediated knockdown of Itch in (**D**) HCT116 and (**E**) HT29 cells.

**Supplementary Figure 2: Impact of Itch knockdown on TRAIL-R2 cell surface expression in a panel of cell lines.**

FACS analysis of cell surface expression of TRAIL-R2 cell surface expression in **(A)** Lim-1215 Ctrl and Itch KD cell lines, **(B)** Colo 320 Ctrl and Itch KD cell lines, **(C)** KM12 Ctrl and Itch KD cell lines, **(D)** HCT116 cells treated with siRNA, **(E)** HT29 cells treated with siRNA. The fluorescence intensity of cell surface staining is represented as percentage of receptor positive cells compared to an IgG isotype control.

## References

1 Shah SS, Kumar S. Adaptors as the regulators of HECT ubiquitin ligases. Cell Death Differ 2021; 28: 455–472.

2 Yin Q, Han T, Fang B, Zhang G, Zhang C, Roberts ER et al. K27-linked ubiquitination of BRAF by ITCH engages cytokine response to maintain MEK-ERK signaling. Nat Commun 2019; 10: 1870.

3 Azakir BA, Desrochers G, Angers A. The ubiquitin ligase Itch mediates the antiapoptotic activity of epidermal growth factor by promoting the ubiquitylation and degradation of the truncated C-terminal portion of Bid. Febs J 2010; 277: 1319–1330.

4 Clorennec CL, Lazrek Y, Dubreuil O, Sampaio C, Larbouret C, Lanotte R et al. ITCH-dependent proteasomal degradation of c-FLIP induced by the anti-HER3 antibody 9F7-F11 promotes DR5/caspase 8-mediated apoptosis of tumor cells. Cell Commun Signal 2019; 17: 106.

5 Chang L, Kamata H, Solinas G, Luo J-L, Maeda S, Venuprasad K et al. The E3 ubiquitin ligase itch couples JNK activation to TNFalpha-induced cell death by inducing c-FLIP(L) turnover. Cell 2006; 124: 601–13.

6 Angers A, Ramjaun AR, McPherson PS. The HECT domain ligase itch ubiquitinates endophilin and localizes to the trans-Golgi network and endosomal system. The Journal of biological chemistry 2004; 279: 11471–11479.

7 Yin Q, Wyatt CJ, Han T, Smalley KSM, Wan L. ITCH as a potential therapeutic target in human cancers. Semin Cancer Biol 2020; 67: 117–130.

8 Roberts JZ, Crawford N, Longley DB. The role of Ubiquitination in Apoptosis and Necroptosis. Cell Death Differ 2021; : 1–13.

9 Lafont E, Hartwig T, Walczak H. Paving TRAIL’s Path with Ubiquitin. Trends Biochem Sci 2018; 43: 44–60.

10 Jarpe MB, Widmann C, Knall C, Schlesinger TK, Gibson S, Yujiri T et al. Anti-apoptotic versus pro-apoptotic signal transduction: Checkpoints and stop signs along the road to death. Oncogene 1998; 17: 1475–1482.

11 Abbas R, Larisch S. Killing by Degradation: Regulation of Apoptosis by the Ubiquitin-Proteasome-System. Cells 2021; 10: 3465.

12 Walczak H, Miller RE, Ariail K, Gliniak B, Griffith TS, Kubin M et al. Tumoricidal activity of tumor necrosis factor–related apoptosis–inducing ligand in vivo. Nat Med 1999; 5: 157–163.

13 Johnstone RW, Frew AJ, Smyth MJ. The TRAIL apoptotic pathway in cancer onset, progression and therapy. Nat Rev Cancer 2008; 8: 782–798.

14 Wang S. The promise of cancer therapeutics targeting the TNF-related apoptosis-inducing ligand and TRAIL receptor pathway. Oncogene 2008; 27: 6207–6215.

15 Özören N, El-Deiry WS. Defining Characteristics of Types I and II Apoptotic Cells in Response to TRAIL. Neoplasia 2002; 4: 551–557.

16 Scaffidi C, Schmitz I, Zha J, Korsmeyer SJ, Krammer PH, Peter ME. Differential Modulation of Apoptosis Sensitivity in CD95 Type I and Type II Cells*. J Biol Chem 1999; 274: 22532–22538.

17 Barretina J, Caponigro G, Stransky N, Venkatesan K, Margolin AA, Kim S et al. The Cancer Cell Line Encyclopedia enables predictive modelling of anticancer drug sensitivity. Nature 2012; 483: 603–607.

18 Ganten TM, Koschny R, Sykora J, Schulze-Bergkamen H, Büchler P, Haas TL et al. Preclinical Differentiation between Apparently Safe and Potentially Hepatotoxic Applications of TRAIL Either Alone or in Combination with Chemotherapeutic Drugs. Clin Cancer Res 2006; 12: 2640–2646.

19 Evergren E, Cobbe N, McMahon HT. Eps15R and clathrin regulate EphB2-mediated cell repulsion. *Traffic (Copenhagen*, Denmark) 2017; 17: 240.

20 Majkut J, Sgobba M, Holohan C, Crawford N, Logan AE, Kerr E et al. Differential affinity of FLIP and procaspase 8 for FADD’s DED binding surfaces regulates DISC assembly. Nat Commun 2014; 5: 3350.

21 Abedini MR, Muller EJ, Brun J, Bergeron R, Gray DA, Tsang BK. Cisplatin Induces p53-Dependent FLICE-Like Inhibitory Protein Ubiquitination in Ovarian Cancer Cells. Cancer Res 2008; 68: 4511–4517.

22 Yang F, Tay KH, Dong L, Thorne RF, Jiang CC, Yang E et al. Cystatin B inhibition of TRAIL-induced apoptosis is associated with the protection of FLIPL from degradation by the E3 ligase itch in human melanoma cells. Cell Death Differ 2010; 17: 1354–1367.

23 Panner A, Crane CA, Weng C, Feletti A, Parsa AT, Pieper RO. A Novel PTEN-Dependent Link to Ubiquitination Controls FLIPS Stability and TRAIL Sensitivity in Glioblastoma Multiforme. Cancer Res 2009; 69: 7911–7916.

24 Shi B, Tran T, Sobkoviak R, Pope RM. Activation-induced Degradation of FLIPL Is Mediated via the Phosphatidylinositol 3-Kinase/Akt Signaling Pathway in Macrophages*. J Biol Chem 2009; 284: 14513–14523.

25 Youn YS, Shin MJ, Chae SY, Jin C-H, Kim TH, Lee KC. Biological and physicochemical evaluation of the conformational stability of tumor necrosis factor-related apoptosis-inducing ligand (TRAIL). Biotechnol Lett 2007; 29: 713–721.

26 Chen J-J, Shen H-CJ, Rosado LAR, Zhang Y, Di X, Zhang B. Mislocalization of death receptors correlates with cellular resistance to their cognate ligands in human breast cancer cells. Oncotarget 2012; 3: 833–842.

27 Jin Z, McDonald ER, Dicker DT, El-Deiry WS. Deficient Tumor Necrosis Factor-related Apoptosis-inducing Ligand (TRAIL) Death Receptor Transport to the Cell Surface in Human Colon Cancer Cells Selected for Resistance to TRAIL-induced Apoptosis*. J Biol Chem 2004; 279: 35829–35839.

28 Azakir BA, Angers A. Reciprocal regulation of the ubiquitin ligase Itch and the epidermal growth factor receptor signaling. Cell Signal 2009; 21: 1326–1336.

29 Artykov AA, Yagolovich AV, Dolgikh DA, Kirpichnikov MP, Trushina DB, Gasparian ME. Death Receptors DR4 and DR5 Undergo Spontaneous and Ligand-Mediated Endocytosis and Recycling Regardless of the Sensitivity of Cancer Cells to TRAIL. Frontiers Cell Dev Biology 2021; 9: 733688.

30 Gonzalvez F, Schug ZT, Houtkooper RH, MacKenzie ED, Brooks DG, Wanders RJA et al. Cardiolipin provides an essential activating platform for caspase-8 on mitochondria. J Cell Biology 2008; 183: 681–696.

31 Lucken-Ardjomande S, Montessuit S, Martinou J-C. Bax activation and stress-induced apoptosis delayed by the accumulation of cholesterol in mitochondrial membranes. Cell Death Differ 2008; 15: 484–493.

32 Tait SWG, Green DR. Mitochondria and cell signalling. J Cell Sci 2012; 125: 807–815.

33 Stöhr R, Mavilio M, Marino A, Casagrande V, Kappel B, Möllmann J et al. ITCH modulates SIRT6 and SREBP2 to influence lipid metabolism and atherosclerosis in ApoE null mice. Sci Rep-uk 2015; 5: 9023.

34 Yu W, Gong J-S, Ko M, Garver WS, Yanagisawa K, Michikawa M. Altered Cholesterol Metabolism in Niemann-Pick Type C1 Mouse Brains Affects Mitochondrial Function*. J Biol Chem 2005; 280: 11731–11739.

35 Dai Y, Zhao X-J, Li F, Yuan Y, Yan D-M, Cao H et al. Truncated Bid Regulates Cisplatin Response via Activation of Mitochondrial Apoptosis Pathway in Ovarian Cancer. Hum Gene Ther 2020; 31: 325–338.

36 Hawash IY, Geahlen RL, Harrison ML, Kesavan KP, Magee AI. The Lck SH3 Domain Negatively Regulates Localization to Lipid Rafts through an Interaction with c-Cbl*. J Biol Chem 2002; 277: 5683–5691.

37 Xu L, Qu X, Zhang Y, Hu X, Yang X, Hou K et al. Oxaliplatin enhances TRAIL-induced apoptosis in gastric cancer cells by CBL-regulated death receptor redistribution in lipid rafts. Febs Lett 2009; 583: 943–948.

38 Chang L, Kamata H, Solinas G, Luo J-L, Maeda S, Venuprasad K et al. The E3 Ubiquitin Ligase Itch Couples JNK Activation to TNFα-induced Cell Death by Inducing c-FLIPL Turnover. Cell 2006; 124: 601–613.

39 Rao DS, Bradley SV, Kumar PD, Hyun TS, Saint-Dic D, Oravecz-Wilson K et al. Altered receptor trafficking in Huntingtin Interacting Protein 1-transformed cells. Cancer Cell 2003; 3: 471–482.

40 Henne WM, Boucrot E, Meinecke M, Evergren E, Vallis Y, Mittal R et al. FCHo proteins are nucleators of clathrin-mediated endocytosis. *Science (New York*, NY) 2010; 328: 1281– 1284.

41 Esposti MD, Ferry G, Masdehors P, Boutin JA, Hickman JA, Dive C. Post-translational Modification of Bid Has Differential Effects on Its Susceptibility to Cleavage by Caspase 8 or Caspase 3*. J Biol Chem 2003; 278: 15749–15757.

42 Breitschopf K, Zeiher AM, Dimmeler S. Ubiquitin-mediated Degradation of the Proapoptotic Active Form of Bid A FUNCTIONAL CONSEQUENCE ON APOPTOSIS INDUCTION*. J Biol Chem 2000; 275: 21648–21652.

43 Tait SWG, Vries E de, Maas C, Keller AM, D’Santos CS, Borst J. Apoptosis induction by Bid requires unconventional ubiquitination and degradation of its N-terminal fragment. J Cell Biology 2007; 179: 1453–1466.

44 Höglinger D, Burgoyne T, Sanchez-Heras E, Hartwig P, Colaco A, Newton J et al. NPC1 regulates ER contacts with endocytic organelles to mediate cholesterol egress. Nat Commun 2019; 10: 4276.

45 Schug ZT, Gonzalvez F, Houtkooper RH, Vaz FM, Gottlieb E. BID is cleaved by caspase-8 within a native complex on the mitochondrial membrane. Cell Death Differ 2011; 18: 538– 548.

46 Akpinar B, Safarikova B, Laukova J, Debnath S, Vaculova AH, Zhivotovsky B et al. Aberrant DR5 transport through disruption of lysosomal function suggests a novel mechanism for receptor activation. Oncotarget 2016; 7: 58286–58301.

47 Montero J, Morales A, Llacuna L, Lluis JM, Terrones O, Basañez G et al. Mitochondrial Cholesterol Contributes to Chemotherapy Resistance in Hepatocellular Carcinoma. Cancer Res 2008; 68: 5246–5256.

48 Wang S-F, Chou Y-C, Mazumder N, Kao F-J, Nagy LD, Guengerich FP et al. 7-Ketocholesterol induces P-glycoprotein through PI3K/mTOR signaling in hepatoma cells. Biochem Pharmacol 2013; 86: 548–560.

49 Wu Y, Si R, Tang H, He Z, Zhu H, Wang L et al. Cholesterol reduces the sensitivity to platinum-based chemotherapy via upregulating ABCG2 in lung adenocarcinoma. Biochem Bioph Res Co 2015; 457: 614–620.

50 Zhong C, Fan L, Li Z, Yao F, Zhao H. SREBP2 is upregulated in esophageal squamous cell carcinoma and co-operates with c-Myc to regulate HMGCR expression. Mol Med Rep 2019; 20: 3003–3010.

51 Kanmalar M, Sani SFA, Kamri NINB, Said NABM, Jamil AHBA, Kuppusamy S et al. Raman spectroscopy biochemical characterisation of bladder cancer cisplatin resistance regulated by FDFT1: a review. Cell Mol Biol Lett 2022; 27: 9.

52 Smith B, Land H. Anticancer Activity of the Cholesterol Exporter ABCA1 Gene. Cell Reports 2012; 2: 580–590.

53 Montero J, Mari M, Colell A, Morales A, Basañez G, Garcia-Ruiz C et al. Cholesterol and peroxidized cardiolipin in mitochondrial membrane properties, permeabilization and cell death. Biochimica Et Biophysica Acta Bba – Bioenergetics 2010; 1797: 1217–1224.

54 Yang Y, Luo M, Zhang K, Zhang J, Gao T, Connell DO et al. Nedd4 ubiquitylates VDAC2/3 to suppress erastin-induced ferroptosis in melanoma. Nat Commun 2020; 11: 433.

55 Baggetto LG, Clottes E, Vial C. Low mitochondrial proton leak due to high membrane cholesterol content and cytosolic creatine kinase as two features of the deviant bioenergetics of Ehrlich and AS30-D tumor cells. Cancer Res 1992; 52: 4935–41.

56 Crain RC, Clark RW, Harvey BE. Role of lipid transfer proteins in the abnormal lipid content of Morris hepatoma mitochondria and microsomes. Cancer Res 1983; 43: 3197– 202.

57 Feo F, Canuto RA, Bertone G, Garcea R, Pani P. Cholesterol and phospholipid composition of mitochondria and microsomes isolated from morris hepatoma 5123 and rat liver. Febs Lett 1973; 33: 229–32.

58 Parlo RA, Coleman PS. Enhanced rate of citrate export from cholesterol-rich hepatoma mitochondria. The truncated Krebs cycle and other metabolic ramifications of mitochondrial membrane cholesterol. J Biol Chem 1984; 259: 9997–10003.

59 Rivel T, Ramseyer C, Yesylevskyy S. The asymmetry of plasma membranes and their cholesterol content influence the uptake of cisplatin. Sci Rep-uk 2019; 9: 5627.

60 Mayengbam SS, Singh A, Pillai AD, Bhat MK. Influence of cholesterol on cancer progression and therapy. Transl Oncol 2021; 14: 101043.

61 Craig EL, Stopsack KH, Evergren E, Penn LZ, Freedland SJ, Hamilton RJ et al. Statins and prostate cancer—hype or hope? The epidemiological perspective. Prostate Cancer P D 2022; : 1–9.

62 Stopsack KH, Gerke TA, Andrén O, Andersson S-O, Giovannucci EL, Mucci LA et al. Cholesterol uptake and regulation in high-grade and lethal prostate cancers. Carcinogenesis 2017; 38: 806–811.

