## Supplementary figures and images for "The E3 ubiquitin ligase Itch regulates death receptor and cholesterol trafficking to affect TRAIL-mediated apoptosis"

### Supplementary Figure 1

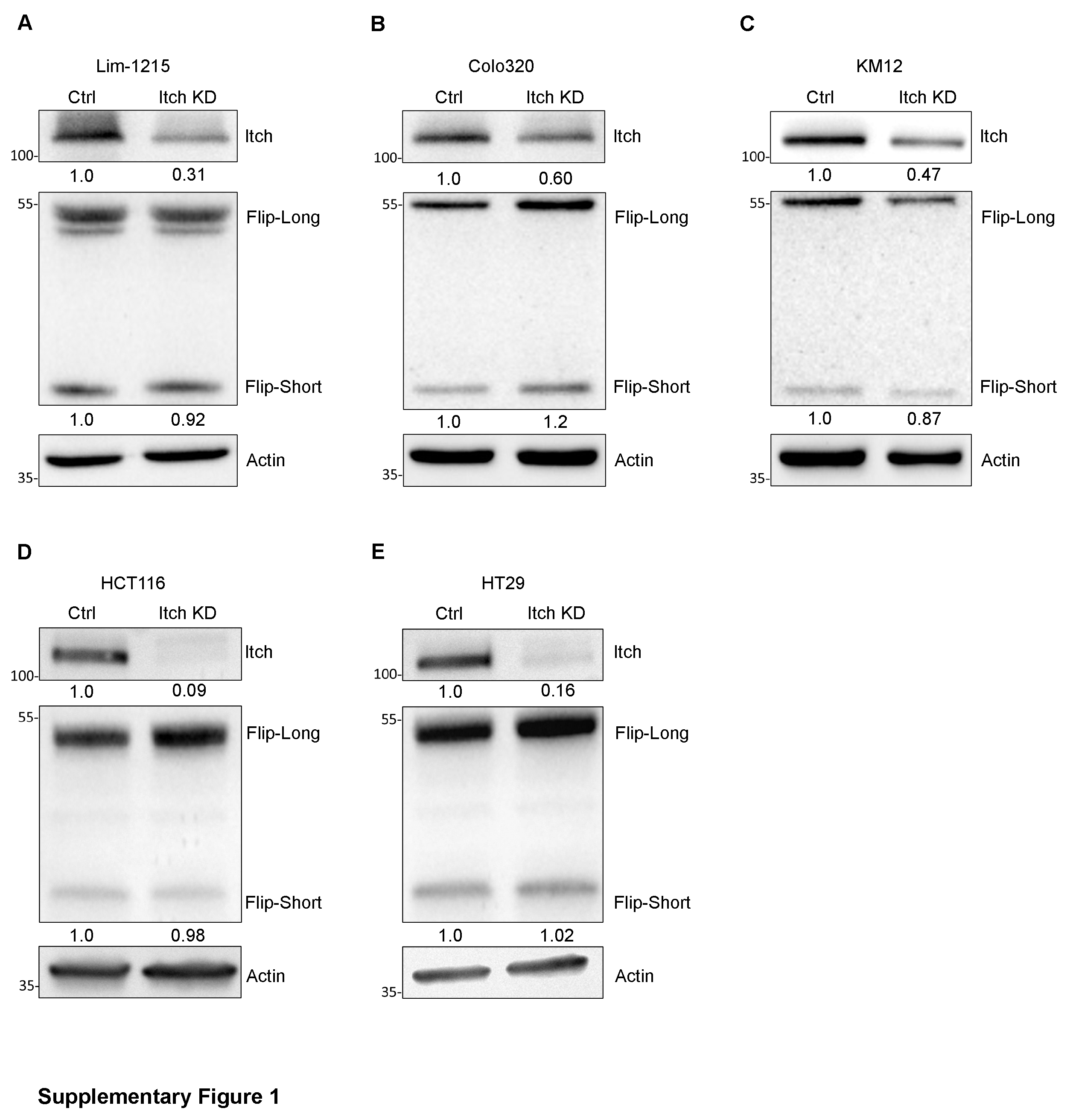

### Supplementary Figure 2

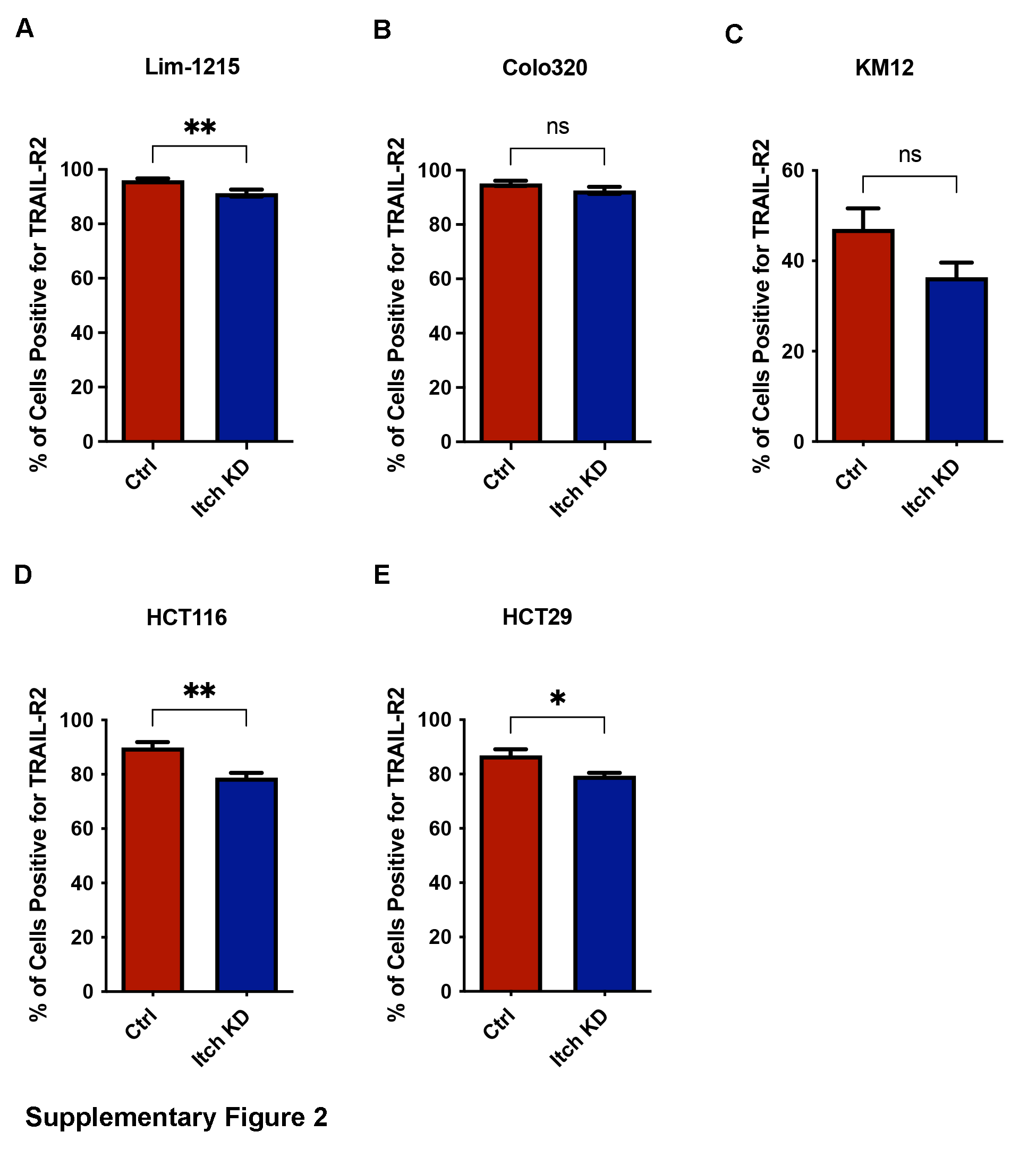
